# Computational tools for serial block EM reveal differences in plasmodesmata distributions and wall environments

**DOI:** 10.1101/2020.03.31.005991

**Authors:** Andrea Paterlini, Ilya Belevich, Eija Jokitalo, Ykä Helariutta

## Abstract

Plasmodesmata are small channels that connect plant cells. While recent technological advances have facilitated the analysis of the ultrastructure of these channels, there are limitations to efficiently addressing their presence over an entire cellular interface. Here, we highlight the value of serial block electron microscopy for this purpose. We developed a computational pipeline to study plasmodesmata distributions and we detect presence/absence of plasmodesmata clusters, pit fields, at the phloem unloading interfaces of *Arabidopsis thaliana* roots. Pit fields can be visualised and quantified. As the wall environment of plasmodesmata is highly specialised we also designed a tool to extract the thickness of the extracellular matrix at and outside plasmodesmata positions. We show and quantify clear wall thinning around plasmodesmata with differences between genotypes, namely in the recently published *plm-2* sphingolipid mutant. Our tools open new avenues for quantitative approaches in the analysis of symplastic trafficking.

**Sentence summary:** We developed computational tools for serial block electron microscopy datasets to extract information on the spatial distribution of plasmodesmata over an entire cellular interface and on the wall environment the plasmodesmata are in.

## Introduction

The cellular units of complex organisms have an intrinsic need for communication. In plants, effective signal exchange is enabled by plasmodesmata (PD), small channels connecting neighbouring plant cells (reviewed in Nicolas *et al*., 2017a). While research has largely focused on the structure and biological regulation of the aperture of the PDs, new insights point to the importance of PD spatial arrangements and their cell wall environments for the flow of materials through them (Deinum *et al*., 2019). Novel methods to get comprehensive information on PD are therefore now required.

The nanometre size of PD pores poses a challenge for their study. A trade-off exists between resolving the detailed structure of the channels and capturing their overall distribution. Electron microscopy (EM) resolved the structure of these channels, identifying a continuous plasma membrane (PM) and a constricted form of the endoplasmic reticulum (ER), the desmotubule, running across the pore between the two cells (Lopez-Saez *et al*., 1966; Robards, 1971). More recently, with the application of electron tomography, variable apposition of PM-ER membranes was shown (Yan *et al*., 2019; Nicolas et al., 2017b). Classical electron microscopy can also be used to study PD occurrence. An inventory of PD densities along the *Arabidopsis* root highlighted interesting variation between cellular interfaces, which might underpin qualitative or quantitative differences in PD-mediated communication between cells (Zhu *et al*., 1998). However, EM approaches, when looking at single or separate slices, largely lose information about the positions of PD relative to each other and only capture approximate densities. This is problematic because distribution of PD across an interface is predicted to have a significant impact on flow properties (Deinum *et al*., 2019). Limited alternatives to comprehensively address the presence of PD have since emerged. Confocal microscopy was applied in leaves, using specific PD markers, to show that the development and distribution of particular PD morphologies in the epidermis was strongly increased by treatments eliciting nutrient, osmotic and pathogen stresses (Fitzgibbon *et al*., 2013). Fluorescent approaches are, however, limited to relatively accessible cell-cell interfaces and often can’t resolve the signal from individual PD. Faulkner *et al*. 2008 used freeze fractured trichomes and EM to analyse PD distributions across the entire fractured surface. They observed that new PD (not generated during cell division) seemed to insert themselves in close proximity to existing PD, suggesting the use of the latter as nucleation centres. The process ultimately results in clusters of PD “pit fields”. A method to obtain similar interface level estimates of PD densities, in this case in the mesophyll layer of leaves, was introduced by Danila *et al*., (2016), combining 3D immuno-localisation, to determine the area of pit field relative to that of the interface, and scanning EM, to assess the number of PD per pit field. However, both immunochemistry-scanning EM and freeze fracture approaches remain confined to tissues that are readily accessible to such sample processing.

Serial block face electron microscopy (SB-EM) (Denk and Horstmann, 2004) can overcome these limitations, offering the opportunity to look at interfaces deep in tissues. A block of fixed and embedded tissue is mounted inside a scanning EM and the upper face of the block is cut away using an internal microtome. After each slice, the newly exposed block surface is imaged. The process is repeated, ultimately generating a stack of images along a z-axis with the z-resolution defined by the thickness of the slices. Importantly, the positions of cellular objects are retained relative to one another and the datasets are good starting points for 3D reconstruction (reviewed in Kittelmann *et al*., 2016). SB-EM technology has been successfully employed to study PD, demonstrating defects in sieve pore (a modified form of PD) structure and distribution (Dettmer *et al*., 2014), and allowing quantification of PD densities at the interfaces of the sieve element (SE) (Ross-Eliott *et al*. 2018) and at the endodermal (EN) face of phloem pole pericycle (PPP) cells (Yan *et al*., 2019). Both SE and PPP cells are key players in the process of phloem unloading, largely mediated by PD (reviewed in Truernit, 2017). These datasets are, however, underexploited in part due to limitations in the technology to extract such information from them. Consequently, important parameters such as the specific distributions of PD, and the cell wall environment of the pores, despite being contained in these datasets, have so far been ignored.

Here we address these two biological aspects. Dense clustering of PD into pit fields is often assumed as a general feature of these structures (Sager and Lee, 2018). However, while this is certainly the case at some interfaces (Danila *et al*., 2016: Faulkner *et al*., 2008), additional evidence is needed to support a generalization. Recent modelling efforts have highlighted how the arrangement of PD in clusters might actually reduce flow between cells (compared to a random arrangement) (Deinum *et al*., 2019). Having detailed information on distributions in actual cells would greatly inform these models. The local wall environment in which PD reside is also of relevance for flow. The thickness of the wall at PD defines the length of the path substances have to travel before entering the neighbouring cell. Thinning at PD is often assumed but the evidence is not comprehensive and quantifications are not available. Correlations between wall thicknesses and different PD ultrastructures of have been reported (Nicolas *et al*., 2017b) and this is now being integrated into models, with predicted effects on flow (Deinum *et al*., 2019). We also know that the PD environment is peculiar in terms of wall polysaccharides, with an enrichment in callose and pectins and a concomitant reduction in cellulose (reviewed in Knox and Benitez-Alfonso, 2014). Overall, the properties generated by wall components have not been extensively explored in planta (partly due to the difficulty of efficiently imaging phenotypic effects).

To extract the relevant information from SB-EM datasets, we developed novel computational and visualization tools dedicated to PD analysis. We deployed the SB-EM datasets from Yan *et al*. (2019) as a study case. We first address the spatial distribution of PD. We detected clusters of PD at the PPP-EN interface while we didn’t see signs of clustering at the SE-PPP interface. We quantified the number and size of the clusters. We quantified specific wall thinning at PD positions and we detected changes in the wall environment in the *plm-2* Arabidopsis mutant.

## Methods

### Datasets

For details on the equipment and settings for SB-EM image acquisition we refer to the original Yan *et al*., 2019 paper, for which the datasets were generated. Briefly, chemically fixed 5 day-old Arabidopsis roots were sectioned and imaged with cutting steps of 40 nm and XY resolutions of 7-10 nm. The collected images were assembled into a single calibrated, aligned and contrast normalised image stack. The resulting images were loaded into the MIB software (Supp. Fig. 1A) (Belevich *et al*., 2016), downloadable at http://mib.helsinki.fi/ (Last accessed March 2020). The images were filtered to reduce noise using deep neural network algorithms, which preserve the edges of the organelles (Supp. Fig. 1B) (Zhang *et al*., 2017). Tutorials on how to operate the software and its tools are available on the website. For the analysis presented here we trimmed the original datasets, each relating to a root and containing multiple cells, into separate datasets for each cell (8 for Col-0 and 5 for *plm-2*). One of the original datasets for the *plm-2* mutant had to be discarded as the image quality was not sufficient for the specific purposes of this study, reducing the available cell number. Removal of the dataset was performed before data analysis.

### Annotations of PD and cell wall segmentations

For PD annotations we re-deployed those from Yan *et al*., 2019. These are contained in the annotation layer of MIB, each annotation including the X, Y, and Z coordinates of the corresponding PD. For analysis, all coordinates were re-calculated from pixels to physical units of the dataset (µm) relative to the bounding box of each dataset. In order to avoid duplicate counts, no new PD were annotated within 160 nm (+2, -2 slices from a central one) of an existing annotation, which provides a conservative estimate (Supp. Fig. 1B). Within the MIB environment, we fully segmented the cell walls of interest by employing black-and-white thresholding within preselected masked areas (Supp. Fig. 1C). Selection of the masked area encapsulating the cell wall was done using the brush tool and an interpolation process to infer the drawn areas on intermediate slices. Resulting models were smoothed and filtered so that the cell wall formed one continuous object in the 3D space. The final model was manually checked for any possible impurities. Small (less than 5 pixels in size) 2D profiles within the 3D model that might not be reliable were removed. High quality segmentations are the basis of any analysis employing the plugins we describe in this paper and they are the most time consuming component of the pipeline.

### Plugins

All the computational tools employed in this paper were written in Matlab language but they don’t require this proprietary software to operate. They are implemented as plugins for the freely available MIB software (Belevich *et al*., 2016). The plugins are included in the standalone version of the MIB software at http://mib.helsinki.fi/downloads.html (Last accessed: March 2020) or separately from https://github.com/AndreaPaterlini/Plasmodesmata_dist_wall (Last accessed: March 2020) In the latter case they need to be saved in the Plugin folder of the MIB software. The plugins are provided with help sections. An overview of the type of file outputs generated by the plugins is provided in Supp.Fig.1. The plugins require as inputs the initial manual steps described in previous sections.

#### The SpatialControlPoints plugin

In order to ask questions relating to the distributions of PD, we felt that a comparable (in terms of points) simulated distribution had to be generated. The simulated distribution differs from the real PD one in that the points are placed with a spatially uniform pattern. To generate such distribution we developed a computational tool, the *SpatialControlsPoints* plugin, capable of creating the “control” point distributions over the same surface as those present in the SB-EM datasets. A segmented wall and a list of annotated PD are fed into the plugin. The tool, in return, finds the midline of the wall by thinning the model to a single centerline without branches (Supp. Fig 1D). The thinning morphological operation (Lam *et al*., 1992) is applied to each slice and then a function detects the longest available path within the thinned lines and removes all the others. The resulting single thin centerline is placed in the mask layer of MIB interface. On this centerline “surface” this tool generates an equal number of points to that of the PD, whose positions are sampled from a uniform distribution, with a randomly placed starting point. For reproducibility of results, in the user interface we provide an option to specify the random seed used by the sampling algorithm. The number of simulated distributions can be defined in the user interface, here we employed 1000 simulations. Matlab and csv file formats are available as outputs (Supp. Fig 1E).

#### The SurfaceArea3D plugin

To calculate the surface of interfaces of interest in the SB-EM datasets we employed an edited and improved version of the plugin used in Yan *et al*., 2019 for the same purpose. The plugin finds the midline of a supplied segmented wall on each slice of the model. This step is the same as that described in the *SpatialControlPoints* plugin (Supp. Fig 1D). This plugin then, additionally, connects such midlines across the slices, generating a surface (Supp. Fig. 1F). The plugin employs the *triangulateCurvePair* function from the *MatGeom* toolbox for geometric computing with Matlab (https://github.com/mattools/matGeom) (Last accessed March 2020). Matlab, Excel and csv file formats are available for the numerical output of the surface. The surface itself can be exported as an object to Matlab, Amira and Imaris programmes.

#### The CellWallThickness plugin

In order to explore the environment surrounding PD, namely the cell wall they span, we developed the *CellWallThickness* plugin, to extract wall thickness from SB-EM datasets (Supp. Fig 1G). The plugin is fed a segmented wall and finds its centerline, as described for the *SpatialControlPoints* plugin (Supp. Fig 1D). A distance map, which assigns a value to each model pixel based on its distance to the closest edge of the model, is calculated at each slice using the Euclidean distance transformation algorithm (Maurer *et al*., 2003)(Supp. Fig. 1H). The values at each point of the masked centerline are then obtained by placing the centerline over the distance map image (Supp. Fig. 1I). The values are expressed in pixels. Since the image is calibrated, the plugin then recomputes those numbers to actual physical thickness of the wall as *thickness (in um) = value (in pixels) x pixel size x 2*. The doubling factor is introduced to obtain wall thickness (and not just half thickness). The masked centerline, where each pixel encodes rounded thickness of the cell wall at the corresponding point can also be saved as an image file. Employing the annotated PD positions, the plugin looks for the closest position on the midline. A line over the wall to show ½ of the distance is displayed (Supp. Fig. 1J). In addition to PD position, if requested, it generates a random uniform distribution over the same surface (employing the *SpatialControlPoints* plugin)(Supp. Fig. 1E) and samples an equal number of points to those of the PD. It can also extract the wall thickness at all points. Depending on the task, the values of real PD and randomly placed PD can be excluded from the list of all points using the corresponding option checkboxes. This ensures independence of classes for statistical comparisons. Matlab, Excel and csv file formats are available as outputs (Supp. Fig 1G).

### R Scripts and guided pipeline availability

The data obtained from MIB and its plugins were then imported and analysed in R (R core team, 2017) to obtain a range of descriptive statistics. We calculated pairwise Euclidean distances between PD (or simulated points) to describe their distribution. The *scatterplot3d* package (Ligges and Machler, 2003 – version 0.3-41) was used in this context to visualise PDs in 3D space. Kolmogorov-Smirnov (KS) tests were used to assess signs of clustering relative to the uniform distributions. Two clustering algorithms were employed to detect the number of clusters present at the PPP-EN interface in root cells. The first is a k-means method with a silhouette approach for estimating optimal cluster number (termed “silhouette” in the figures), which was implemented using the *factoextra* package (Kassambara and Mundt, 2017 – version 1.0.5). The second one is a Bayesian Information Criterion for expectation-maximization, initialized by hierarchical clustering for parameterized Gaussian mixture models (termed “mclust”), which was implemented using the *mclust* package (Scrucca *et al*., 2016 – version 5.4.2). In both cases we arbitrarily defined the maximum numbers of clusters to 20, believing the 1:20 range to be biologically meaningful. In the case of the k-means algorithm we additionally repeated the initial seed placing 100 times, in order to reduce the possibility of inaccurate clustering due to biases in initial seed placement. To determine the surface areas occupied by the identified clusters we projected the 3D coordinates of the PD onto a 2D space using principal component analysis (PCA). No significant loss of information in the distribution of the PD occurred (likely relating to the fact that the cell walls were mostly flat planes). The two first principal components of PCS captured >90% of the variance in the x-y-z coordinates of the original data in all cases reported here. The areas of the convex hulls delimited by the outer points of each cluster (or the outer points in general, in the case of the total surface) were extracted using the *splancs* package (Rowlington and Diggle, 2017 – version 2.01-40). Lastly, we estimated the cell wall thickness around PD. A guided tutorial with all the necessary code for this analysis is available at https://andreapaterlini.github.io/Plasmodesmata_dist_wall/ (Last accessed March 2020). The Col-0 datasets used in this paper, with corresponding models and annotation are available from https://drive.google.com/file/d/1g-Wg7AuJTwVdNdB3ydkkET8hvQtFOPC0/view **(/these will be deposited in an official data repository before final submission**), and can be used as an example dataset to follow our tutorial. In addition to the specific packages listed above we also employed the broader *tidyverse* environment (Wickham, 2017 – version 1.3.0) and the *data*.*table* (Dowle and Srinivasan, 2019 – version 1.12.0) and *ggbeeswarm* (Clarke and Sherrill-Mix, 2017 – version 0.6.0) packages.

### 3D visualisations

For 3D visualisation we employed both Imaris (version 8.4.2) and Amira (version 2019.1) imaging software. Export of features from the MIB environment are compatible with both visualization packages.

## Results

### SB-EM allows spatial positioning of PD over wall interfaces

SB-EM datasets can cover large portions of a tissue. The datasets from Yan *et al*., 2019 employed here cover an area encompassing the cells around the protophloem of Arabidopsis roots. The datasets can be visualised either in a longitudinal orientation (Fig 1A) or in an axial one (Fig 1B). In the latter, PD at various radial interfaces are more easily detectable due to better XY-resolution of the SB-EM technique (Fig 1C and D showing PD at SE-PPP and PPP-EN interfaces respectively). The image resolution of the specific datasets employed here is good enough to identify individual PD within those areas, but not to distinguish the detailed morphology of the PD. A unique aspect of SB-EM is that such annotated PD positions (relative to the cell surface) can be addressed globally within the full length of cells. While density calculation approaches have taken advantage of this (Yan *et al*., 2019), the spatial component, namely the 3D distribution of PD, has been largely neglected. Traditional bi-dimensional visualisations fail to convey the distribution of PD over the interfaces. Here we show that identified PD can be exported (as clouds of dots) alongside the segmented wall, generating effective 3D spatial representations that capture the distribution. We show this both at the SE-PPP and PPP-EN interfaces (Fig 1E and 1F). The rendering can also be stored as movies (SuppMovie1).

**Figure 1.**
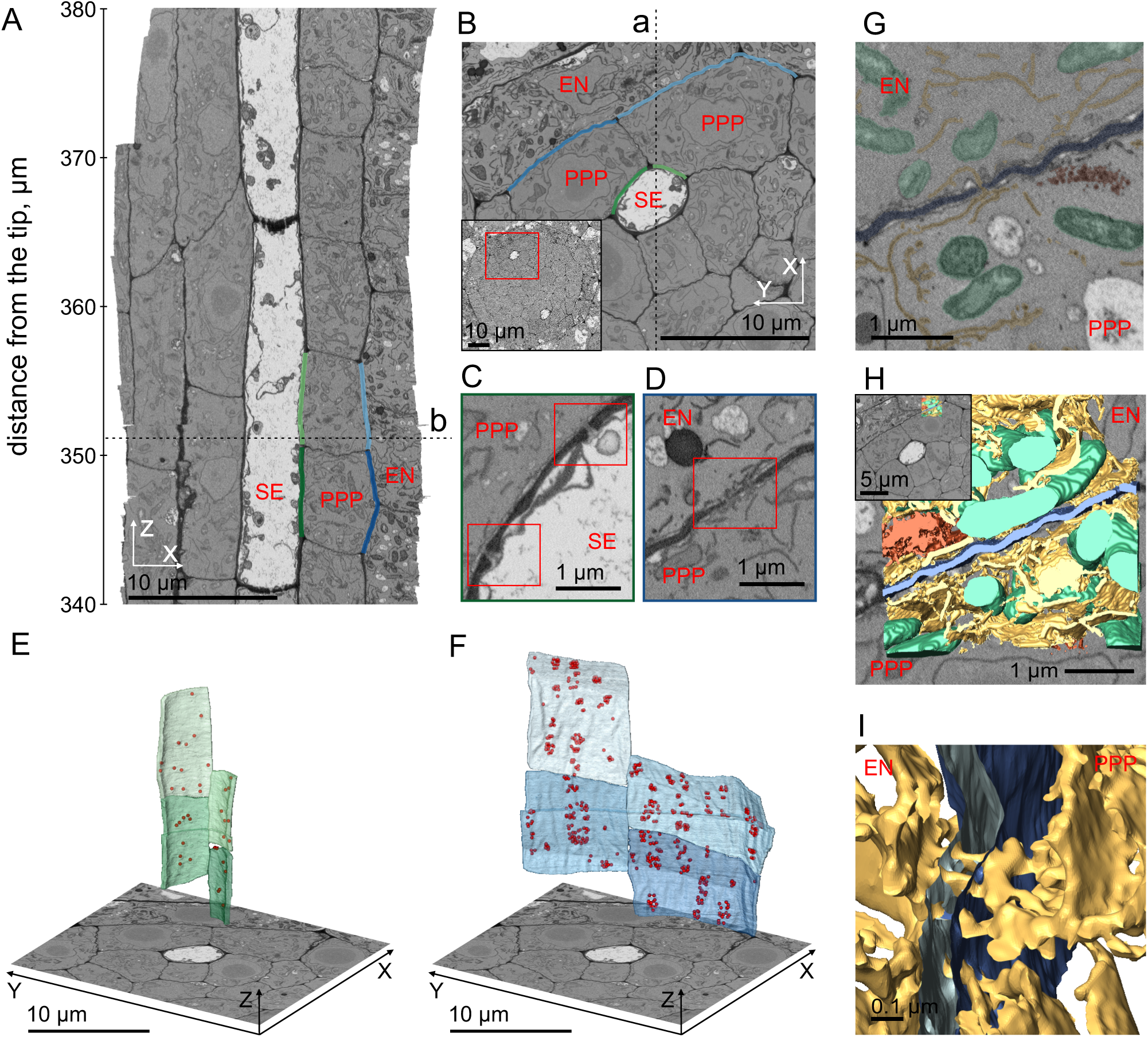
Overview of SB-EM technology and its spatial capabilities on Col-0 datasets. A) Example dataset slice seen in a xz orientation. The sieve element (SE) is in the centre. Its interface with the phloem pole pericycle (PPP) is shown in green and the interface between the PPP and endodermis (EN) to is in blue. The dashed line marked by b shows the relationship to the view in B). B) Extract of The same dataset visualised in xy orientation. The inset panel displays the acquired area relative to an overview of the root. The dashed line marked by a shows the relationship to the view in A). C)D) Zoomed areas of the SE-PPP and PPP-EN interfaces respectively showing PD within the red rectangles. E)F) 3D visualisation in the Amira software of SE-PPP and PPP-EN interfaces in four cells (colour coded green or blue as in A). The wall is segmented and PD are represented as red dots. G) Two dimensional view of an area along the PPP-EN interface with organelles and structures highlighted (ER in yellow, mitochondria in green, Golgi in red, wall in blue). H) 3D visualisation in the Amira software of the segmented organelles and structures from G. The inset panel displays the position of the model (top right) relative to the area of interest in B. I) Zoomed in area of the model shown in H, seen from the sides, showing ER strands crossing the PPP-EN wall.

### Rendering of PD in the cellular context

SB-EM datasets also contain data on various organelles within the cells, putting PD in a wider and more realistic context. We can generate highly structured and dense cellular models. By segmenting an area of the datasets (represented in the overall dataset in Fig 1H, inset panel) and colour coding the different organelles (Fig 1G) the 3D model can be eventually exported in visualization programmes (Fig 1H). We can show the ER strands of PD crossing into the wall (Fig 1I) and then merging on either sides to the wider ER network system. Various animations can then be applied to the 3D model (SuppMovie 2). Models such as these highlight how symplastic transport needs to navigate a dense cytoplasm before reaching the PD.

### Signs of clustering of PD at the PPP-EN interface in the root

Taking advantage of the spatial information on PD contained in the SB-EM datasets, we studied their distribution at selected interfaces. To describe the distribution of PD on the cell wall, we calculated pairwise Euclidean distances between each of them, using the x,y,z coordinates available in the datasets. This revealed a multi-modal distribution of distances (red lines in Figure 2A, where we show two examples of cells). To provide a meaningful comparison an equal number of points with a uniform distribution was generated on the same surface using the *SpatialControlsPoints* tool described in the methods. One thousand simulations were generated for each cell. We calculated Euclidean distances for each of the simulations and observed that in each case the distributions of distances approximated a normal distributions (yellow lines in Figure 2A, for the two example cells). The surfaces are not perfectly flat, resulting in deviations from full normality. This immediately suggests some overall differences. Next to each plot for the distribution of distances we also show the original 3D distribution of points, to give an appreciation of the biases in point distribution between real data and simulation (Fig 2B, showing the same two example cells). At the PPP-EN interface, in addition, an excess of short distances between PD points was always visually detectable compared to the distributions of distances of the simulated points. This suggests some form of clustering (e.g. Fig 2A). For each of the 8 cells tested for Col-0 or the 5 cells tested for *plm-2* (the mutant available in the Yan *et al*., 2019 datasets) pairwise KS tests of the distribution of real points against each one of the 1000 simulated point sets were performed. The distribution of p-values is shown in Fig C (in pink for PPP-EN interface). All cells fell below a p-value of 0.05 (black vertical line), suggesting a non-uniform distribution of PD. To give a quantitative appreciation of variation across cells in spatial distributions, the distribution of the KS test result values can be plotted (higher KS test values relate to stronger differences between the real and simulated distributions) (Fig 2D). Overall, test values for the single comparisons ranged from 0.025 to 0.347. Figure 2A and B actually displayed the two most extreme cases among cells of the Col-0 genotype: we took the cell with the distribution most shifted to lower KS values (PPP1-ENa) and that shifted to higher values (PPP2a-EN), these are shaded in a darker grey in the figure. The spatial plots in Fig 2B match well the expectations. Comparisons between genotypes can be performed by plotting mean KS test values for each cell of both genotypes (Fig 2E). Summary values per cell remove the otherwise present problem of interdependencies of points, which would complicate statistical comparisons. The *plm-2* genotype did not show appreciable shifts compared to WT (medians of 0.09 and 0.08 respectively).

**Figure 2.**
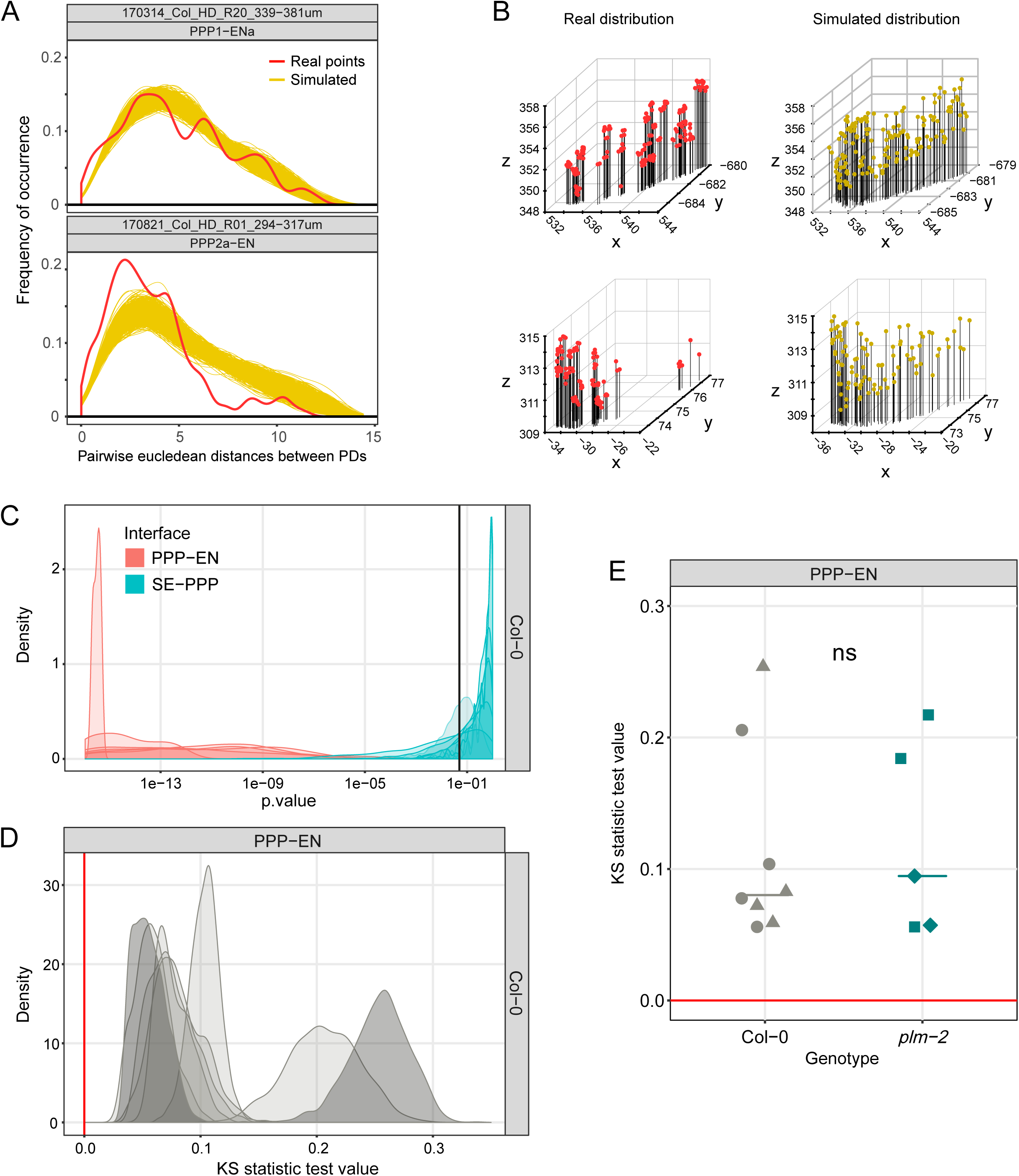
Detection of spatial clustering at cellular interfaces using SB-EM datasets. A) Distribution of Euclidean distances between points at the PPP-EN interface in two cells (top and bottom panels). The red line represents distances between PD while each of the 1000 yellow lines represents the distances between uniformly distributed control points (in each simulation). B) 3D visualisation of PD positions (red, left panels) and the uniformly distributed control positions in one of the simulations (yellow, right panels) for the two cells shown in A) (top and bottom panels). C) Distribution of p-values of KS test at the PPP-EN or SE-PPP interfaces for the Col-0 cells (8000 p-values for each interface: 8 cells x 1000 simulations each). Black vertical bar highlights the 0.05 value, used as a significance threshold. Distribution of KS test values at the PPP-EN interface for the Col-0 cells (8000 p-values in total). Red bar highlights the 0 value, representing identity between real and simulated distributions. Left dark grey curve - top panel cell in (A), right dark grey curve - bottom panel cell in (A). E) Comparison of KS test values at the PPP-EN interface between Col-0 and *plm-2* genotypes. Values for single cells (8 for Col-0 and 5 for *plm-2*) are represented with symbols (different symbols for different roots), medians as horizontal bars. Statistical comparisons between genotypes were performed using the non-parametric Mann-Whitney U test for two samples, ns=p>0.05.

### Lack of PD clustering at the SE-PPP interface in the root

At the SE-PPP interface in Col-0, conversely, there was no evidence to reject the null hypothesis of a uniform distribution for the PD. The p-values for the KS test comparison between real and simulated points were shifted towards or above 0.05 in Fig 2C (light blue curves). Mean p-values for each cell were above 0.05. We show the distribution of Euclidean distances for one of the cells. The distribution of distances for the real points appeared less diverse relative to of the simulated points, compared to those observed at PPP-EN interface. In addition, no visual excesses of shorter distances could be detected (Supp.Fig. 2A). The 3D distributions of real and simulated points are shown in Supp.Fig. 2B. The distributions of the KS test values, while being at times higher than those at PPP-EN interface in terms of absolute values, were much shallower (Supp.Fig.2C, the dark shaded cell in this panel has been used as the example in Supp. Fig. 2A,B).

Because the SE-PPP interface has a lower number of PD compared to the PPP-EN interface (Supp.Fig. 2D) we tried to assess if the high p-values at the SE-PPP interface were just due to lower statistical test power or were an indication of actual lack of clustering. We sampled a lower number of PDs (and simulated points) at the PPP-EN to achieve the same PD density as that seen at the SE-PPP interface. The number of new points was calculated by multiplying the density of PDs at the SE-PPP interface by the surface area at the PPP-EN interface (Supp.Fig. 2D). We then tested if a difference between the distribution of Euclidean distances of real points and simulated ones could still be detected. While the p-values did indeed on average shift towards higher values, in 6/8 cells of Col-0 the mean p-value was still below 0.05 (red vertical line) (Supp.Fig. 2E). In only two cells (purple ones in the figure) the PD distribution could no longer be robustly differentiated from a uniform one. Overall, we feel this suggests that at the SE-PPP interface there are indeed no obvious signs of PD clustering and this highlights differences between this interface and the PPP-EN interface.

### Describing the organisation of PD in pit fields at the PPP-EN interface

Upon establishing the presence of a non-uniform distribution of PD over the PPP-EN interface we attempted to characterise the potential clusters. Namely, we tried to address the number of clusters, the number of PD per cluster and the cluster sizes relative to the surface of the interface. To determine the number of clusters, we used two different clustering algorithms, a k-mean based method and a model based one, within the R environment. Variation was visible - as should be expected due to the relatively arbitrary computational classification - between the single cells and between clustering algorithms with median values of 11.5 (Col-0) and 10 (*plm-2*) clusters using the mclust package and 11.5 (Col-0) and 14 (*plm-2*) in the silhouette approach (Fig 3A). Differences between genotypes were not statistically significant so, overall, a working range of 10-14 PD clusters can be suggested at the PPP-EN interface. As an example we colour coded the PD of a cell according to the cluster they had been assigned with two methods (Fig 3B and 3C). In the image the 3D coordinates had been reduced to 2D via PCA. Some of the strengths and pitfalls of these clustering methods are illustrated in this example, with the silhouette approach being possibly over-conservative while mclust assigned 1 PD to the wrong cluster (the olive green dot in the bright green cluster in Fig 3B, highlighted with an asterisk). We strongly emphasize that cluster number values should be used as working ranges rather than absolute values. The number of PD in each cluster was similar between the two clustering methods, with a median of 8-10 PD/cluster (Fig 3D). Once again, no strong trends in the *plm*-2 mutant from Yan *et al*., 2019 datasets were detectable.

**Figure 3.**
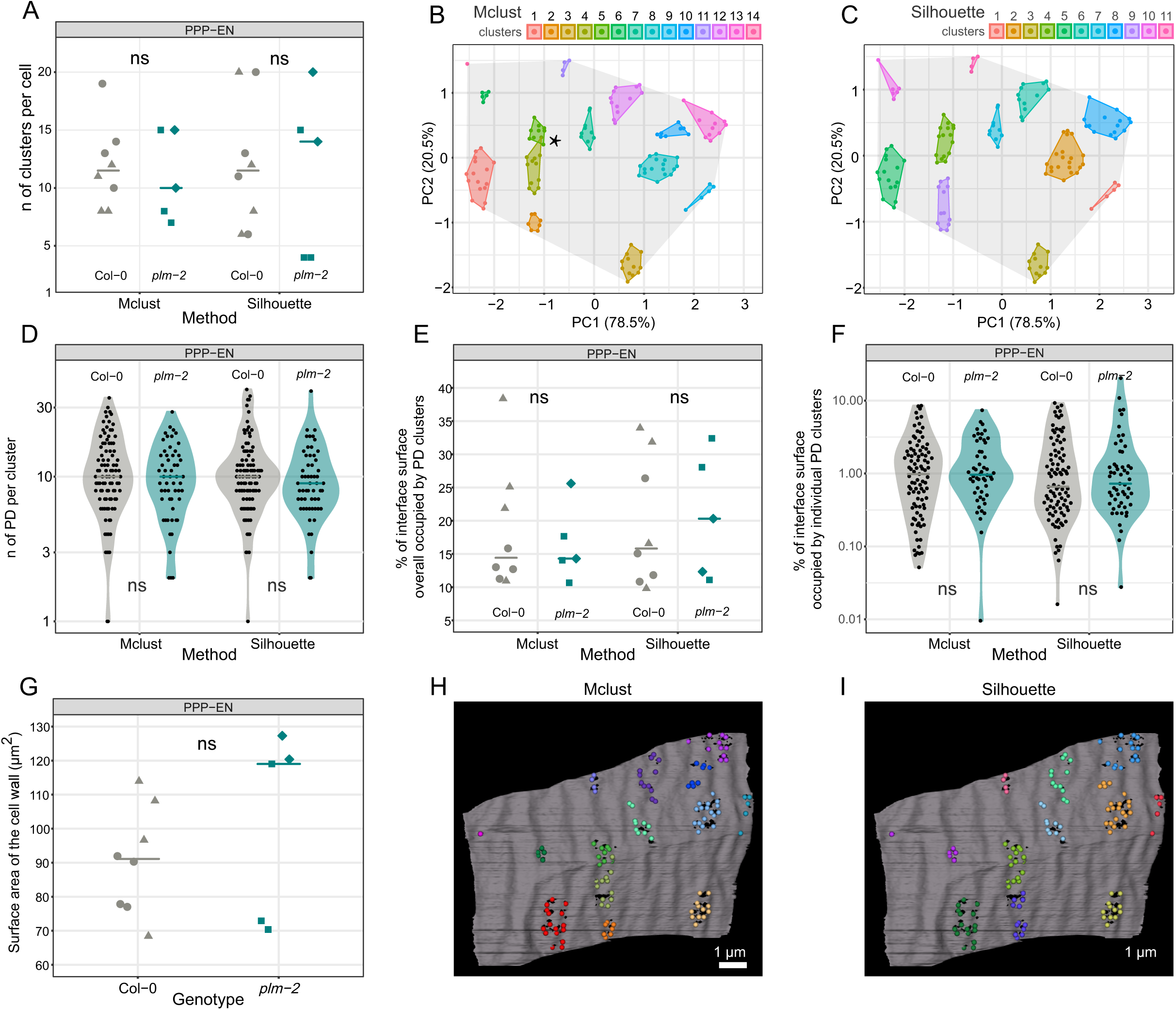
Quantification of clustering parameters at the PPP-EN interface using SB-EM datasets. A) Number of clusters identified in the Col-0 (8 cells) and *plm-2* (5 cells) genotypes using the mclust or the silhouette approaches. B)C) Visualisations of a PCA reduced interface (from a cell) with different cluster assignments. The surface is rendered in grey while PD belonging to different clusters and the area they occupy are colour coded. D) Number of PD per cluster in Col-0 and *plm-2* genotypes. The total clusters for Col-0 are 95 (Mclust) and 96 (Silhouette) while 52 (Mclust) 55 (Silhouette) for *plm-2*. Total % of the surface occupied by clusters. F) % of the surface occupied by individual clusters. G) Absolute surface in µm^2^ of cells. H)I) 3D visualisations in the Imaris programme of the same interface shown in B)C). The surface is rendered in grey while PD belonging to different clusters are colour coded with the same scheme used previously. Please note that D and F have logarithmic y-axes. In the graphs, individual values are represented as dots, distributions as violin plots, and medians as horizontal bars. Cells from different roots are shown using different symbols. For each clustering approach, statistical comparisons between genotypes were made using the non-parametric Mann-Whitney U test for two samples, ns=p>0.05.

To assess the “conductive” surfaces provided by these PD clusters, we calculated the area occupied by each of these clusters and that occupied in total by all the clusters on a cell interface. These areas were calculated and reported as percentages of the total surface of the interface on which they occur, as delimited by the most extreme clusters points in the PCA space (shaded grey area in Fig 3B and C). While this is an underestimate of the total surface we feel it is more appropriate to calculate these scaling factors rather than attempting to use the cluster surfaces as absolute values (the data are indeed scaled and reduced by the PCA so the units are no longer true µm^2^). For the total “conductive” surface, i.e. the proportion occupied by the clusters relative to the overall surface, the mclust method suggests medians of 14% for both Col-0 and *plm-2* while the Silhouette approach provides median values of 15% and 18% respectively (Fig 3E). Each cluster accounts for a median surface of 1% in both genotypes using Mclust method or 0.7% (Col-0) and 0.8% *(plm-2*) with the silhouette approach (Fig 3F). No differences between the two genotypes were highlighted by statistical testing for any of these parameters.

While these might be sufficient for some purposes we also wanted the possibility to relate these percentages to the actual surface values in µm^2^. To do so, we had to employ the original image data within the MIB software, rather than the R processed and PCA reduced ones. Using an updated version of the *SurfaceArea3D* plugin employed in Yan *et al*., 2019 we calculated the actual total surface of the PPP-EN interfaces in MIB (see methods for details). A median surface area of 91.1 µm^2^ was determined for Col-0 and 119 µm^2^ for *plm-2* (Fig 3G). Given the variance in the data, this difference is not robust. Relating the median surface area values to the scaling factors described above we can obtain the actual surfaces of the individual clusters. The surface can be exported to Imaris for visualization (Fig 3H and 3I, mirroring Fig 3B and 3C).

### Extracting and visualising wall thickness at PD, controls and every (other) position

The SB-EM datasets contain information of many components of a cell, in the context of PD a relevant one is the cell wall and its thickness. To do that we developed the *CellWallThickness* plugin, that can extract the thickness of a given segmented wall at positions of interest (see methods section). As an example, we employ the plugin on one of the Col-0 cells available (the one used in Fig 3B and C). By using an equal number of “random uniformly distributed points” (median of 117 nm) one can accurately capture the thickness of the overall “all other” wall (median of 123 nm) in a computational effective manner. These values are not statistically significantly different. The data also show a clearly thinner wall in the proximity of “PD” (median of 46 nm)(Fig 4A). The thickness of the wall at the interface of interest can also be visualised graphically, by exporting the midline thickness map (generated by the plugin) into 3D rendering software such as Imaris (Fig 4B). The wall colour intensity matches the calculated thickness value at that position, brighter meaning thicker. The PD positions and those of the controls can also be exported as dots and their relative size made to match the wall thickness value. The thinning at PD positions is visually confirmed and shown to extend beyond the precise position of the channels, to the entire pit field PD are grouped into.

**Figure 4.**
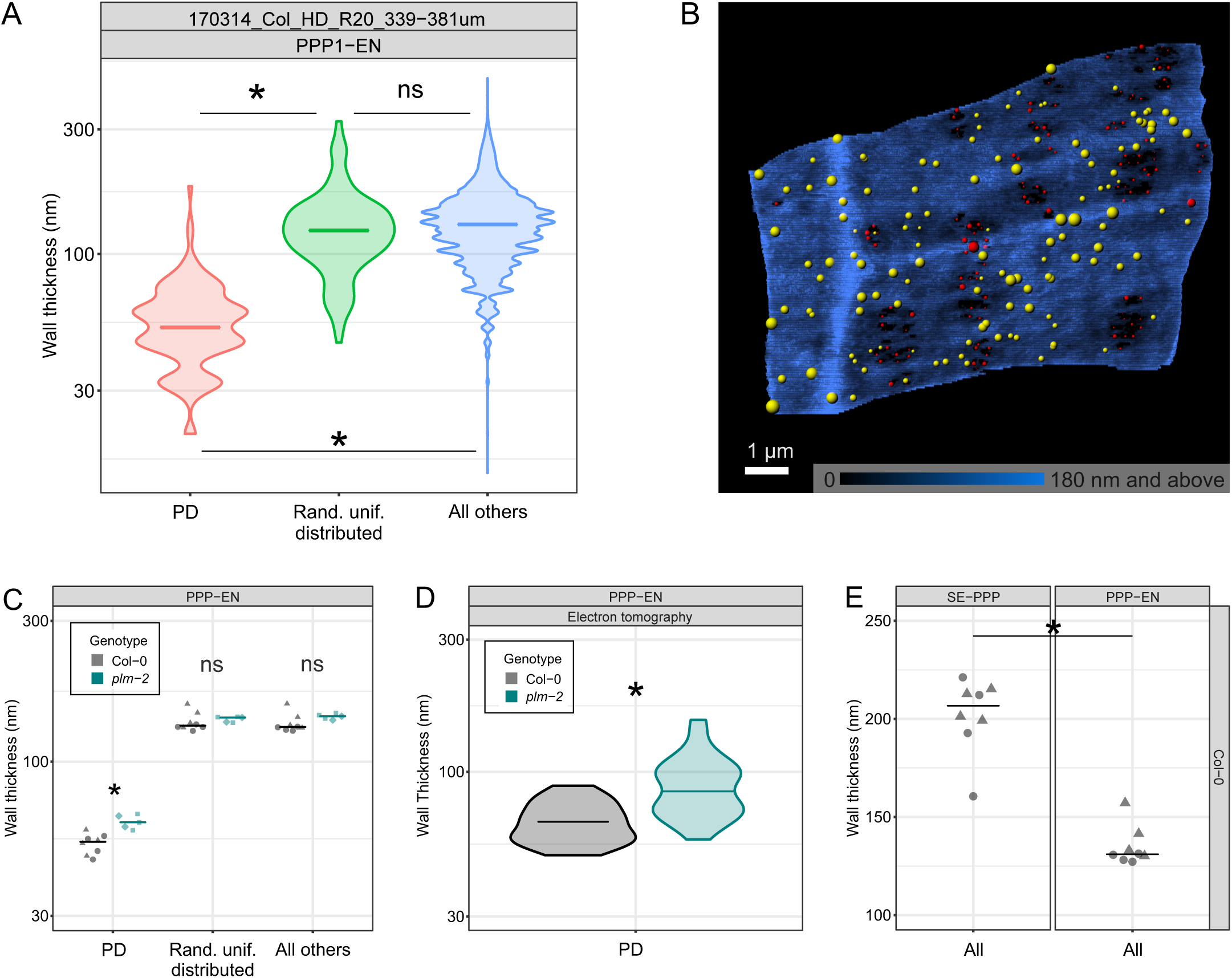
Assessments of cell wall thickness using SB-EM datasets. A) Wall thickness for the PPP-EN interface in one Col-0 cell at PD positions, random uniform control positions and at all other points (n=129 for PD positions and random uniform control positions, n=222864 for all points). B) 3D visualisation in the Imaris programme of a PPP-EN interface in Col-0, as in Fig 3. The surface is rendered in shades of blue, on a scale matching the thickness of the wall. PD are shown as coloured dots (red for real PD positions, yellow for those of a simulation with a random uniform distribution). The size of the dots relates to the wall thickness at that position. C) Comparison of wall thickness in Col-0 (8 cells) and *plm-2* (5 cells) genotypes. D) Comparison of wall thickness at PD positions in Col-0 and *plm-2* genotypes using tomography data (n=30 for Col-0 and n=49 for *plm-2*). E) Comparison of overall wall thickness at the SE-PPP and PPP-EN interfaces in Col-0 (8 cells). In the graphs average values for single cells are represented with symbols (different symbols for different roots), distributions as violin plots, medians as horizontal bars. Please note that A, C and D have logarithmic y-axes. Statistical comparisons between two samples (or two samples within a category) were performed using the non parametric Mann-Whitney U test, or for more than two samples the non-parametric Dunn’s test. Supported differences are highlighted by an * (p<0.05).

Cell wall thickness comparisons between genotypes are also possible with our tools, using mean thickness values for the three categories of points in each cell. At PD positions the median thickness of the resulting values was 53 nm for Col-0 and of 62 nm for *plm-2* (Figure 4C). The difference is supported by statistical testing. Conversely, no difference was supported for the “random uniformly distributed points” (132 nm in Col-0 vs 141 nm in *plm-2*) nor for the “all other points” category (131 nm in Col-0 vs 142 nm in *plm-2*), despite a trend for increased thickness in *plm-2* (Figure 4C).

To test if the difference at PD positions between genotypes could be independently confirmed, we looked at another dataset contained in the Yan *et al*., (2019) paper. Electron tomography, a different technique, had been employed to study the ultrastructure of single PD at the PPP-EN interface in the two genotypes. When we re-deployed those data, this time extracting the wall thicknesses in the immediate proximity of PD, we obtained median values of 66 nm in Col-0 and 85 nm for the *plm-2* mutant (Fig 4D). Statistical testing supported a difference. These values are remarkably close to those obtained with our plugin: an absolute match was unlikely considering the difference in scale of observation (individual PD compared to the entire tissue). Both techniques therefore agree in showing a trend of thicker walls at PD (and possibly across the entire wall) in the *plm-2* mutant.

To further validate the reliability of the data generated we assessed if known biological features could be detected in our datasets and if the values obtained matched those from different techniques. The wall of enucleated SEs is thicker compared to that of nucleated SEs or of surrounding cells. This reinforcement is likely necessary to withstand the pressure of sap flow (Furuta *et al*., 2014). The plugin output was able to effectively capture and quantify this difference using the “all points” category. The median thickness of the SE-PPP wall, using averages per single cells, was of 207 nm compared to 131 nm for the PPP (Fig 4E). Note that here “all points” are used rather than “all other points” as there is no need to exclude PD positions.

## Discussion

PD perform a key role in cell-cell transport across plant cells. We explored aspects of their distributions and of the wall they span. We performed this in part with the aim of informing future models of flow across PD with relevant experimental data. Published modelling approaches have so far studied PD transport in relation to the overall single pore structure (Blake, 1978), phloem flow (Jensen *et al*., 2012), phloem loading mechanisms (Comtet *et al*., 2017) and unloading flow type (Ross-Elliott *et al*., 2017). One modelling study tried to address some complexities of PD, integrating ultrastructure parameters for the cytoplasmic sleeve in their models (Liesche and Schultz 2013). The assumptions of this paper, however, have been challenged by experimental data (Ding *et al*., 1992; Nicolas *et al*., 2017b), highlighting the difficulty of modelling flow across PD when limited experimental data are available. Only recently, the spatial distribution of the PD at interfaces, namely the assumption of clustering in pit fields, is starting to be included in models. Unequal distribution was shown to reduce the effective symplastic permeability of the interface (Deinum *et al*., 2019). However, detailed experimental data for distribution parameters, such as those we present here, is lacking. Including PD spatial arrangements in models is a significant advancement from the use of the same spatial parameters (number of clusters and PD per cluster) to estimate the total conductive surface alone. Using traditional microscopy approaches Kuo *et al*., 1974 had for instance calculated assimilate fluxes in wheat leaves and, more recently and more comprehensively, Danila *et al*., 2016 had used immunolabelling in accessible leaf tissues to compare fluxes in C3 and C4 monocot leaves. However, such studies might have underestimated the impediment to flow such PD arrangements may impose.

We show that at an interface important for post-phloem unloading in the Arabidopsis root, that of PPP-EN (Yan *et al*., 2019), there is clear spatial clustering of PD (Fig 2A, B, C). We determine and visualise the numbers of pit fields at the PPP-EN interface, with a median of 10-14 clusters (Fig 3A). Computer driven clustering methods present their own limitations in terms of absolute value generation, possibly explaining part of the observed variation within and between clustering approaches. However, pit field numbers in different cells might also be affected by the surface and age of the actual cell. This in turn would explain the clustering metric variation observed in Fig2 A. We don’t have enough cells spanning the z direction to empirically test such a hypothesis but it remains plausible and interesting for future research. Overall, we recommend using these values as working ranges rather than absolute values. The remarkably stable median value for the number of PD per pit fields, around 10 with all methods (Fig 3D) might conversely hint at some biological process eventually constraining cells to create new clusters as they expand. There might be an upper boundary for secondary PD formation within a cluster. Lastly, in cells the orientation of the PD and the overarching clusters (Fig 1F or 3H,I) often seem to follow the direction of vertical cell elongation.

We also calculated the median surface area of the clusters and related it to the total surface (Fig 3E, G). An approximate median conductive surface of 15% (each cluster accounting for a median of around 1%) is a remarkably low percentage and certainly suggests the possibility that flow in and out of a cell might not be uniform, but rather resembles more a series of water currents in a cytoplasmic ocean. It will be extremely interesting to apply such concept to modelling studies of flow between cells as started in Deinum *et al*., 2019.

In our data, we don’t observe clustering at the SE-PPP interface (Fig 2C and Supp.Fig. 2B,C), which is fundamental for phloem unloading in the root (Ross-Eliott *et al*., 2017). A more uniform distribution of PD might reflect the enucleated nature of SEs (Furuta *et al*., 2014) and the impact this might have on secondary PD formation or a unique feature of the funnel PD at this interface, known to perform batch unloading (Ross-Eliott *et al*., 2017). It is interesting that in Deinum *et al*., 2019 clustering of PD had the largest negative effect on parameters regulating flow at lower PD densities. The lower density of PD at the SE-PPP interface compared to neighbouring tissues (Yan *et al*., 2019) might impose limits to clustering if this was to compromise the extensive flow that needs to take place at this interface. Overall, this result at least challenges the broad assumptions that all PD might be grouped into pit fields. The distribution tools presented in this paper could be used for more systematic studies within tissues.

In addition to the spatial distribution of PD, parameters describing the environment surrounding PD can also be of high value. We address wall thicknesses, affecting flow between cells in relation to the structure of PD in recent modelling studies (Deinum *et al*., 2019), with our new tool. The overall wall thickness is in the same order of magnitude as the estimation in Kramer *et al*. (2007). The value of around 200±30 nm was actually derived from a figure in Andème-Onzighi et al. (2002), focused on root epidermal cells. We detect a thinner wall around PD at the PPP-EN interface in the root, by a factor of about 2.5 times (Fig 4C), matching assumptions in the literature that PD lie in wall depressions. The agreement of wall thickness values at PD between SB-EM and electron tomography (a technique that focuses on the area of one PD) is extremely satisfactory although we still caution on using these values as absolute. Thinner walls at PD might be the consequence of cell wall modifications required for PD de-novo insertion (Faulkner *et al*., 2008) or a pre-requisite for PD insertion at all. Regardless of the ontological reason of this wall thinning (or lack of thickening), it is likely that it carries functional relevance for conductivity (Thompson and Holbrook, 2003; Baratt *et al*., 2011). A few reports from other plant species mention that the sieve pores (highly modified forms of PD) in mature plates connecting SEs lie in wall depressions. This was correlated to callose deposition inhibiting wall thickening (Evert *et al*., 1966; Deshpande, 1974, 1975). Whether that is a shared mechanism to all PD, also rich in callose, is unknown. In modelling studies a relative arbitrary value of 100 nm is employed as the wall thickness for PD (Liesche and Schultz, 2013). This is compatible with the averages for our PPP-EN cells (Fig 4C, D). However, our workflow might be highly valuable in future studies to inform cell-cell permeability models of differences between interfaces. For instance, we quantify the known biological difference of SE wall thickness (Furuta *et al*., 2014) compared to PPP cells (Fig 4E).

Our tool could easily be applied to broad cell wall questions. For instance, to our knowledge, in the literature there aren’t tissue specific studies of cell wall thickness in the Arabidopsis root. In addition, although SB-EM is not yet high throughput enough to allow mutant screens, targeted validation of mutants can be performed. We detected thicker walls around PD (and possibly globally) in the *plm-2* mutant compared to WT (Fig 4C,D). The mutant is defective in the biosynthesis of VLCFA sphingolipids (Yan *et al*. 2019). GIPCs sphingolipids, known to be enriched at PD (Grison *et al*. 2015), are cross-linked via boron bridges with pectins (Voxeur *et al*. 2014), also likely enriched at PD (reviewed in Knox and Benitez-Alfonso, 2014). It is therefore reasonable to speculate that there may be feedback effects on wall structure from lipid perturbations. The detected thicker wall emphasises the importance of PD type change observed in this mutant for flow: it must provide a significant ease of trafficking to achieve the reported increase in communication (Yan *et al*., 2019). Alternatively, modelling studies suggest that different types of PD might be more suitable in different types of walls (Deinum *et al*., 2019). Overall, interesting new lines of research might develop from a more systematic use of SB-EM and associated analysis tools.

## Acknowledgements

We would like to thank Hugo Tavares (Sainsbury Laboratory, Cambridge) for his valuable input, support on R-coding aspects of the project and reading of the manuscript. We thank David Legland (Institut National de la Recherche Agronomique, France) for help with triangulation of points. We would also like to acknowledge William Nicolas and Emmanuelle Bayer (Laboratory of Membrane Biogenesis, Bordeaux) for allowing us to use the tomography data we collected in our previous collaboration and for comments on the manuscript. We thank Ottoline Leyser (Sainsbury Laboratory, Cambridge) and Clelia Bolpagni for providing comments to the manuscript.

This work was supported by the Gatsby Foundation (GAT3395/PR3), Biocenter Finland, the Finnish Centre of Excellence in Molecular Biology of Primary Producers (Academy of Finland CoE program 2014-2019, decision #271832), University of Helsinki (award 799992091) and the European Research Council Advanced Investigator Grant SYMDEV (No. 323052).

## Figure legends

*Supp.Video1: Video of the dataset*

*Supp.Video2: Video of the cellular model*

**Supp.Fig.1:**
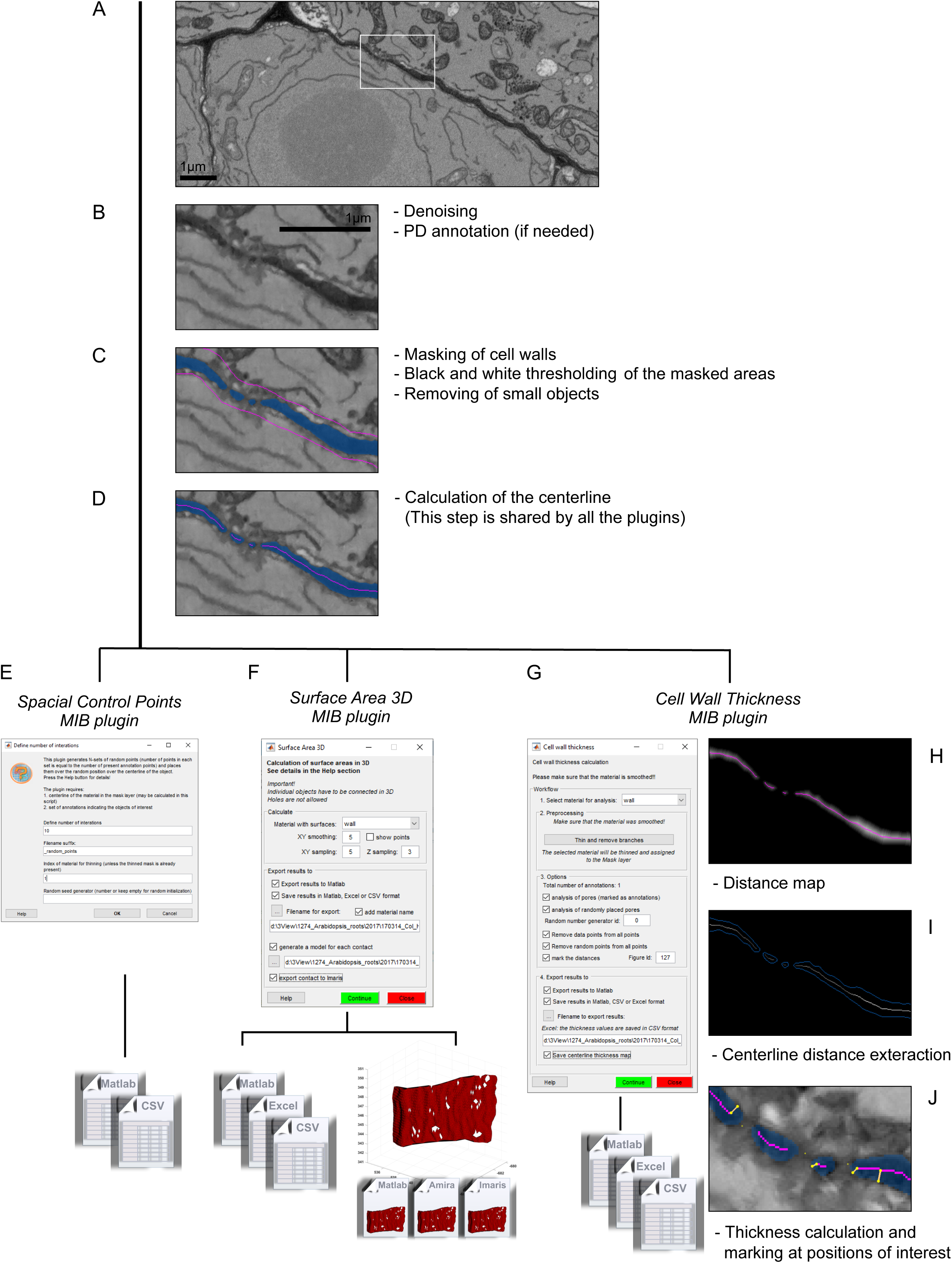
Overview of the plugins developed for this paper. A) Overall view of PPP-EN interface. B) Zoomed in view of the area highlighted in A) after denoising. C) Segmented wall in blue and the underlying mask in pink. D) Calculated midline of the cell wall model. E) User interface and file outputs for the *SpatialControlPoints* plugin. F) User interface, file and object outputs for the *SurfaceArea3D* plugin. G) User interface and file outputs for the *CellWallThickness* plugin. H) Distance map of the segmented wall. I) Thickness values of the midline. J) Yellow lines showing the wall position (and associated half thickness) closest to PD/control points being measured.

**Supp.Fig.2:**
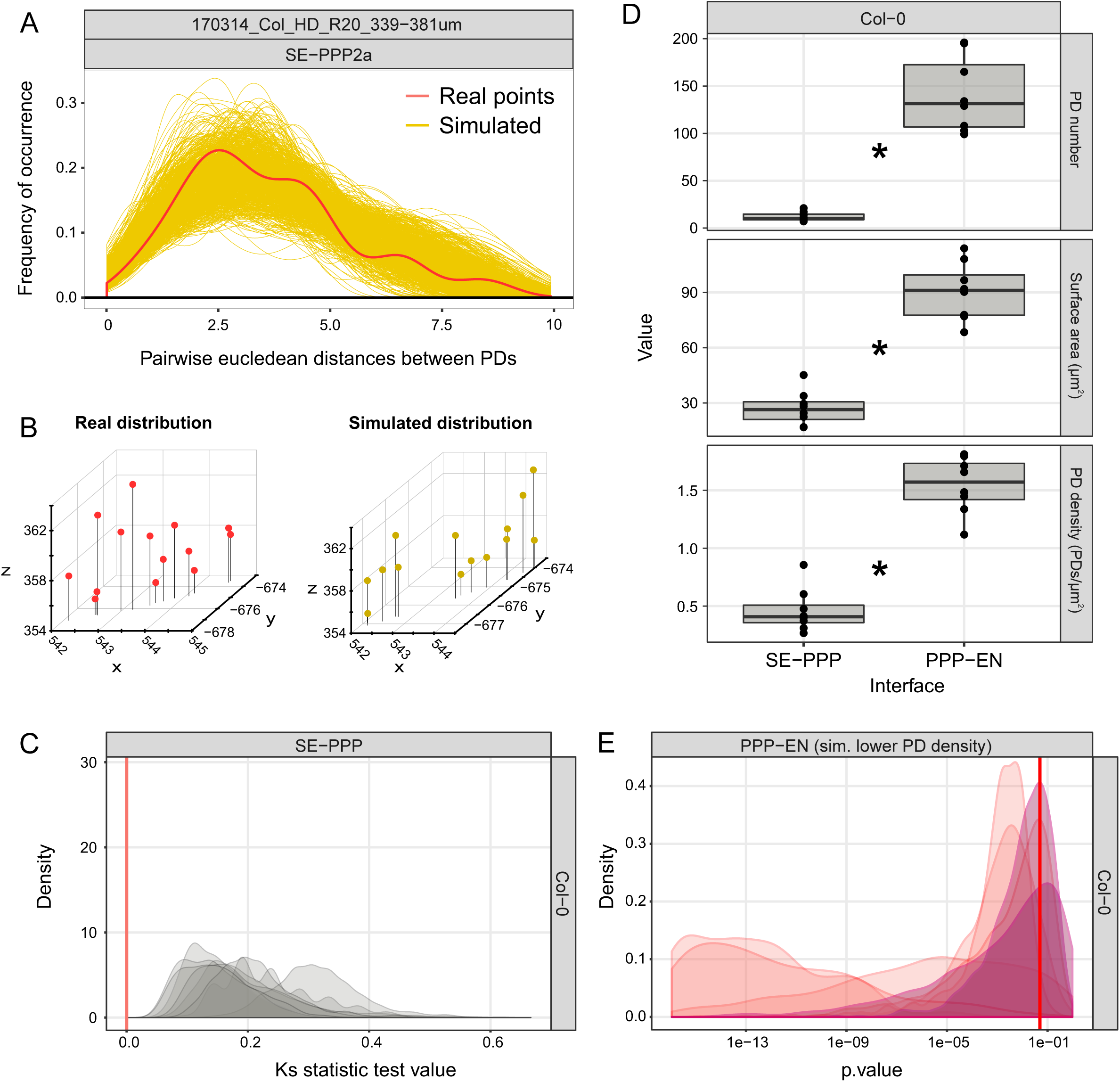
Analysis of the SE-PPP interface in terms of spatial clustering using SB-EM datasets. A) Distribution of Euclidean distances between two points at the SE-PPP interface in one example cell. The red line represents distances between PD while each of the yellow lines represents the distances between uniformly distributed control points (in each simulation). B) 3D visualisation of PD positions (red) and the uniformly distributed control positions in one of the simulations (yellow) for the cell. C) Distribution of KS test values for the Col-0 cells. Red bar highlights the 0 value, representing identity between real and simulated distributions. D) Comparison of PD numbers, surface areas and PD densities at the SE-PPP and PPP-EN interfaces. E) Distribution of p-values of KS test at the PPP-EN interface when using a number of PD matching the density at the SE-PPP interface. Red bar highlights the 0.05 value, used as a significance threshold. Cells shaded have mean p-values above 0.05. Statistical comparisons between genotypes were performed using the non parametric Mann-Whitney U test for two samples. Supported differences are highlighted by an * (p<0.05).

https://drive.google.com/file/d/1P-kdn1RgIydgHrzM_n7RY_37_jeqiMAV/view?usp=sharing

Make sure to increase the video quality in the player setting when viewing it

**Supp. Video 1**

https://drive.google.com/file/d/1I7NAFrs7EE3mqSPbHjxY1VGWMyWXPN3f/view?usp=sharing

Make sure to increase the video quality in the player setting when viewing it

**Supp. Video 2**

